# Probing the aggregated effects of purifying selection per individual on 1,380 medical phenotypes in the UK biobank

**DOI:** 10.1101/2020.11.16.385724

**Authors:** Ha My T. Vy, Daniel M. Jordan, Daniel J. Balick, Ron Do

**Affiliations:** The Charles Bronfman Institute for Personalized Medicine, Icahn School of Medicine at Mount Sinai, New York, NY, USA; Department of Genetics and Genomic Sciences, Icahn School of Medicine at Mount Sinai, New York, NY, USA; Division of Genetics, Brigham and Women’s Hospital, Harvard Medical School, Boston, MA, USA; Department of Biomedical Informatics, Harvard Medical School, Boston, MA, USA

## Abstract

Understanding the relationship between natural selection and phenotypic variation has been a long-standing challenge in human population genetics. With the emergence of biobank-scale datasets, along with new statistical metrics to approximate strength of purifying selection at the variant level, it is now possible to correlate a proxy of individual relative fitness with a range of medical phenotypes. We calculated a per-individual deleterious load score by summing the total number of derived alleles per individual after incorporating a weight that approximates strength of purifying selection. We assessed four methods for the weight, including GERP, phyloP, CADD, and fitcons. By quantitatively tracking each of these scores with the site frequency spectrum, we identified phyloP as the most appropriate weight. The phyloP-weighted load score was then calculated across 15,129,142 variants in 335,161 individuals from the UK Biobank and tested for association on 1,380 medical phenotypes. After accounting for multiple test correction, we observed a strong association of the load score amongst coding sites only on 27 traits including body mass, adiposity and metabolic rate. We further observed that the association signals were driven by common variants (derived allele frequency > 5%) with high phyloP score (phyloP > 2). Finally, through permutation analyses, we showed that the load score amongst coding sites had an excess of nominally significant associations on many medical phenotypes. These results suggest a broad impact of deleterious load on medical phenotypes and highlight the deleterious load score as a tool to disentangle the complex relationship between natural selection and medical phenotypes.

**Author summary:** This study aims to augment our understanding between the complex relation between natural selection and human phenotypic variation. We developed a load score to approximate the relative fitness of an individual and correlate it with a set of medical phenotypes. Association tests between the load score amongst coding sites and 1,380 phenotypes in a sample of 335,161 individuals from the UK Biobank showed a strong association with 27 traits including body mass, adiposity and metabolic rate. Furthermore, an excess of nominal associations at suggestive levels was observed between the load score amongst coding sites and medical phenotypes than would be expected under a null model. These results suggest that the aggregate effect of deleterious mutations as measured by the load score has a broad effect on human phenotypes.

## Introduction

One of the primary questions of interest in the study of human population genetics is the relation between natural selection and the evolution of human phenotypes, from quantitative traits to complex disease. With the emergence of biobank-scale datasets, along with new statistical metrics to approximate strength of purifying selection at the variant level, it is now possible to both estimate the net impact of deleterious mutations for each individual in a large population sample and correlate it to a range of medical phenotypes exhibited by that individual. This provides an opportunity to simultaneously study the genetics of individuals within a relatively homogenous population and the potential impact of natural selection on annotated phenotypes.

A large body of literature exists on evolution and estimation of the deleterious mutation load from human population samples, with particular emphasis on cross-ancestry comparisons [1-6]. Rather than a comparison between human populations, we aimed to assess the distribution of deleterious loads—the sum of all purifying selective effects in each individual’s genome—within a single human population. While the mutation load generally represents a population-wide average of this quantity, we estimated the same object for each individual in the population to produce a “load score” that counts the net effect of deleterious variation in each individual’s genome, a count of derived alleles weighted by an estimate of the selective disadvantage for each variant. When compared to the mean of the population, this per-individual load score can be interpreted as a component of the relative fitness of each individual.

In this study, we aim to augment our understanding of the relation between natural selection and human phenotypes by focusing on the net impact of purifying selection on the fitness of each individual, and correlating this quantity to the set of phenotypes acting on that individual. Previously this has been difficult for two reasons: first, we do not have a direct measure of the fitness of individual humans that can be estimated from genetic information, and second, there were no large databases available to quantify the wide range of phenotypes possessed by each individual. Biobank-scale datasets that contain both individual genotypes and phenotypes, such as the UK Biobank [7, 8], finally provides access to both large-scale phenotypic descriptions of each individual and some part of their genetic sequence.

We ventured to apply computational tools that predict aspects of purifying selection for individual alleles to published genotypes of 335,161 white British individuals from the UK Biobank to estimate the fitness impact of derived variation present in each imputed genome in this sample. Most of the variation in the sample exists at appreciable frequencies, and is likely under relatively small selective disadvantage, but in aggregate the fitness impact can be substantial. Using this representation of each individual’s relative fitness, we probed correlations between the impact of common deleterious variation in an individual’s genome and their personal phenotypic makeup. This provides a different lens into questions about the relation between fitness vis-à-vis mutation load and human traits by looking at a per-individual measure correlated to fitness, rather than focusing on the distribution of selective effects in the population as a whole. This allows us to ask which phenotypes, if any, are highly correlated to the aggregation of deleterious variation, and probe the relation between the ensemble of phenotypes and fitness loss due to common variation in individuals.

## Results

### Comparison of four deleteriousness prediction scoring methods

The additive effects of deleterious variation can be quantified in aggregate by a genome-wide score representing the net action of purifying selection on an individual under the assumption that effects of individual variants can be summed additively. Multiple methods have been developed to characterize purifying selection, including methods that predict deleterious selection acting on the level of a single allele (fitCons [9], FATHMM-MKL [10], deltaSVM [11], Funseq2 [12]), methods that measure evolutionary conservation (phyloP [13], phastCons [14], GERP++ [15], SiPhy [16]) and methods that predict the effect of an allele on molecular function (CADD [17], DANN [18], GenoCanyon [19], Eigen and EigenPC [20]). Although these scores are formulated as tests for strong selection or for molecular function, rather than as estimates of the strength of selection, they are also correlated to the strength of selection, and are often used as proxies for strength of selection [4, 21, 22]. In this study, we compared the predicted deleteriousness of alleles for four widely used scoring methods that approximate deleteriousness of a variant--GERP++, phyloP, CADD, and fitCons--with their effects on allelic frequency in human population genetic data to select the most appropriate measure for computation of the additive load.

Under negative or purifying selection, natural selection acts to reduce the population frequency of deleterious mutations. This effect is more able to overcome genetic drift as the strength of selection increases. As a result, we expect to observe a higher number of rare alleles and a lower number of common alleles in regions of the genome that are under negative selection, relative to putatively neutral regions. This can be seen as a shift of the allele frequency spectrum (AFS) towards rare alleles, with a steeper slope of the AFS indicating stronger purifying selection. We evaluated the extent to which each scoring method captures the deleteriousness of an allele by grouping alleles by the scores provided by each method and measuring the slope of the resulting AFS. The more strongly a score is related to the strength of selection, the more marked the increase in slope will be for high-scoring alleles relative to lower scoring alleles.

We evaluated this correlation using whole genome sequencing data from a non-Finnish European population in the Genome Aggregation Database (gnomAD) [23]. For each scoring method, we grouped alleles by score and compared the non-normalized derived allele frequency spectra for each group (see Methods). Fig 1 plots the log of the derived allele frequency (DAF), and shows a consistent pattern across all scores: the higher the score, the steeper the slope of the log DAF (S1 Table). This indicates that, for all scores shown, higher scores are associated with sites under stronger negative selection, as expected. While all four scores show this pattern, CADD and phyloP show clearer separations between DAF spectra than fitCons and GERP++. In the case of fitCons, this underperformance is likely due to its incorporation of functional genomic signatures that may increase its performance at identifying functional regions, but detract from its performance at identifying sites under purifying selection. In the case of GERP++, the underperformance is more surprising, since GERP++ and phyloP are very similar methods. The difference in performance may be explained by differences in how the final scores computed by the two methods are defined, or by the fact that the scores were calculated using two different multiple sequence alignments: the phyloP scores were calculated from UCSC’s alignment of 100 vertebrate sequences generated using the MultiZ method [24], while the GERP++ scores were calculated from the Ensembl alignment of 111 mammalian sequences generated using the EPO-Extended method [25]. For comparison, we also calculated DAF spectra for synonymous sites, missense sites, and loss of function sites (LOF). Based on this comparison, phyloP scores between 5 and 7.5 appear to be similarly deleterious to missense sites (nearly the same DAF slope) and phyloP scores greater than 7.5 appear to be under similar purifying selection to LOF sites. The equivalent numbers for CADD are 20 to 25 for missense and greater than 30 for LOF.

**Figure 1.**
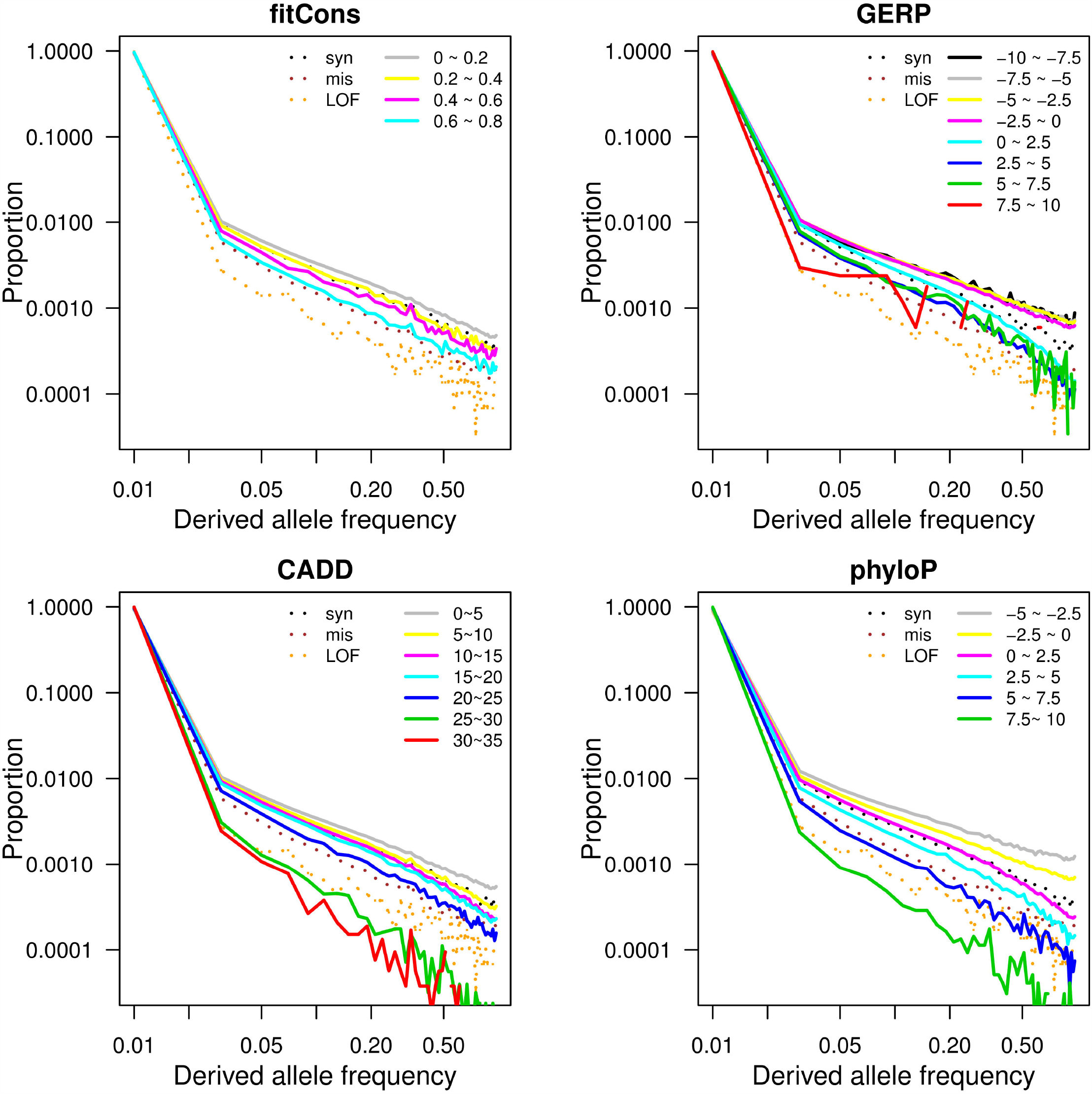
Derived allele frequency spectra of different score categories for each deleteriousness prediction scoring method. For each scoring method, polymorphic sites are grouped into score intervals by the value of the score annotated at the sites. Each solid line represents derived allele frequency spectrum of polymorphic sites belonging to one score interval and three dashed lines represent derived allele frequency spectra of three control categories: synonymous (syn), missense (mis), and loss of function (LOF) variants.

To further compare between CADD and phyloP, we examined the DAF distribution for protein coding variants and noncoding variants separately. Both scores performed similarly for coding variants, but phyloP showed better separation in noncoding variants (S1 Fig). This is as expected, since CADD uses more features when scoring coding variants than noncoding variants, while the phyloP method is identical for coding and noncoding sites. For this reason, we concluded that phyloP has the most consistent relationship between score and strength of negative selection, and selected phyloP as our weight for our load score computation.

### Per-individual load scores in UK Biobank

We calculated a per-individual deleterious load score by summing the total number of derived alleles per individual, weighting each derived allele by its phyloP score to account for the strength of purifying selection. We considered three load scores: a genome-wide load score, a coding-specific load score, and a non-coding-specific load score. Each score was computed for 335,161 unrelated, white-British ancestry individuals in the UK Biobank using 6,774,062 variants from imputed genotypes (95,850 coding and 6,678,212 non-coding) with positive phyloP scores (positive scores denote uniform purifying selection, while negative scores denote clade-specific selection). The observed population distribution across all sampled individuals appear very close to normal for each of our three scores (Kolmogorov-Smirnov Tests P-values = 0.32; 0.55; 0.20 for all variants, non-coding, and coding, respectively, Fig 2). This is the expected result if the phyloP scores of derived alleles are identically distributed across the entire population, due to the Central Limit Theorem. By contrast, if the white-British population contained distinct subpopulations with dramatically different distributions of phyloP scores among derived alleles, we would expect to see a sum of multiple normal distributions with different means, resulting in a skewed or multi-modal distribution.

**Figure 2.**
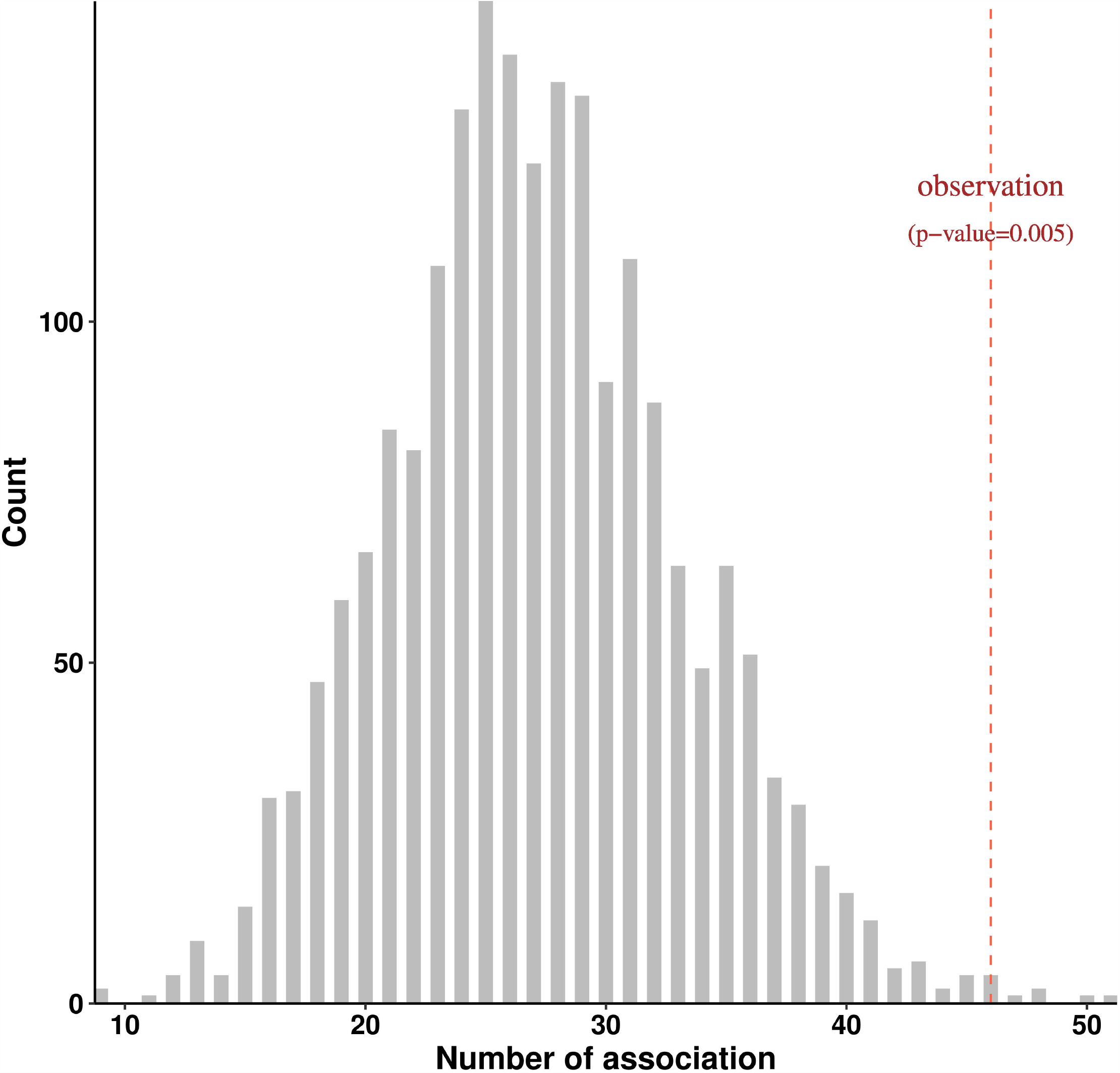
Distribution of load score. Histogram of three load scores computed from three sets of variants: coding variants (coding load score), non-coding variants (non-coding load score), and both coding and non-coding variants (genome-wide load score). Each load score was computed for 335,161 unrelated, white-British ancestry individuals.

### Significant association between load score of coding variants and anthropometric and metabolic traits

To explore the overall effect of deleterious mutations on specific clinically measured phenotypes, we tested the association of each of the three load scores (genome-wide, coding and non-coding) with 1,380 traits, after adjusting for age, sex, genotyping chip, and assessment center. To account for potential confounders, we further included a set of geographical and socioeconomic variables available in the UK Biobank data as additional covariates (S2 Table). We note that many of these variables are significantly associated with the load score but the effects are small. Nonetheless, careful consideration was taken to add these as covariates in our association tests (S2 Table).

We discovered no phenotype significantly associated with either the genome-wide load score or non-coding load score (Bonferroni P value threshold = 1.2×10^−5^). However, 27 traits were significantly associated with the load score calculated from coding SNPs; these included body mass, metabolic rate, and several adiposity traits such as body mass index and waist circumference (Table 1). Some of these traits have been found under directional selection in contemporary populations [26-28].

**Table 1.** Association between coding load score and 27 traits.

| Trait | Sample size | beta | SE | P-value |
| --- | --- | --- | --- | --- |
| Arm fat-free mass (right) | 329238 | -0.0073 | 0.0012 | $6.61 \times 10^{-10}$ |
| Arm fat-free mass (left) | 329182 | -0.0074 | 0.0012 | $7.93 \times 10^{-10}$ |
| Arm predicted mass (left) | 329170 | -0.0073 | 0.0012 | $1.12 \times 10^{-9}$ |
| Leg fat mass (left) | 329295 | -0.0090 | 0.0015 | $1.39 \times 10^{-9}$ |
| Basal metabolic rate | 329326 | -0.0074 | 0.0012 | $2.83 \times 10^{-9}$ |
| Arm predicted mass (right) | 329235 | -0.0069 | 0.0012 | $3.76 \times 10^{-9}$ |
| Weight | 334221 | -0.0098 | 0.0017 | $3.98 \times 10^{-9}$ |
| Whole body water mass | 329333 | -0.0068 | 0.0012 | $8.68 \times 10^{-9}$ |
| Leg fat mass (right) | 329311 | -0.0086 | 0.0015 | $1.02 \times 10^{-8}$ |
| Whole body fat-free mass | 329306 | -0.0067 | 0.0012 | $1.24 \times 10^{-8}$ |
| Whole body fat mass | 328780 | -0.0103 | 0.0018 | $1.80 \times 10^{-8}$ |
| Leg fat percentage (left) | 329297 | -0.0064 | 0.0011 | $2.66 \times 10^{-8}$ |
| Trunk fat-free mass | 329057 | -0.0065 | 0.0012 | $3.55 \times 10^{-8}$ |
| Trunk predicted mass | 329019 | -0.0064 | 0.0012 | $5.07 \times 10^{-8}$ |
| Trunk fat mass | 329118 | -0.0100 | 0.0019 | $1.21 \times 10^{-7}$ |
| Leg predicted mass (right) | 329303 | -0.0063 | 0.0012 | $1.54 \times 10^{-7}$ |
| Leg fat-free mass (right) | 329303 | -0.0063 | 0.0012 | $1.96 \times 10^{-7}$ |
| Leg fat-free mass (left) | 329280 | -0.0062 | 0.0012 | $3.18 \times 10^{-7}$ |
| Leg predicted mass (left) | 329275 | -0.0062 | 0.0012 | $3.77 \times 10^{-7}$ |
| Leg fat percentage (right) | 329316 | -0.0058 | 0.0012 | $5.74 \times 10^{-7}$ |
| Arm fat mass (right) | 329242 | -0.0092 | 0.0018 | $6.38 \times 10^{-7}$ |
| Body mass index (BMI) | 334097 | -0.0092 | 0.0019 | $7.25 \times 10^{-7}$ |
| Arm fat mass (left) | 329188 | -0.0090 | 0.0018 | $1.15 \times 10^{-6}$ |
| Waist circumference | 334612 | -0.0080 | 0.0016 | $1.35 \times 10^{-6}$ |
| Body fat percentage | 329134 | -0.0066 | 0.0014 | $2.98 \times 10^{-6}$ |
| Hip circumference | 334579 | -0.0088 | 0.0019 | $3.29 \times 10^{-6}$ |
| Impedance of arm (left) | 329313 | 0.0061 | 0.0013 | $5.25 \times 10^{-6}$ |

SE: standard error

Stratification by derived allele frequency showed that these association signals are more pronounced when limiting to variants that are common (DAF > 5%) but not close to fixation (DAF < 70%), while stratification by phyloP score shows that they are more pronounced when limiting to variants with higher phyloP scores (phyloP>2, S3 and S4 Tables). We therefore performed an additional stratification analysis by both DAF and phyloP score (S5 Table). We observed that the signals are mostly driven by common variants (5<=DAF<70%) with higher phyloP score (phyloP>2). This class of variants notably contributes a large fraction (mean: 0.38 and sd: 0.005 per individual) towards the per individual coding load score. This analysis necessarily excludes extremely rare alleles, which are not well captured by the process of genotyping and imputation. It is not clear how significant the aggregate contribution of these alleles to the per individual load score would be.

To assess the effect of our weighting procedure, we calculated an unweighted load score, the per-individual mutation burden, that simply counts derived alleles with no reference to phyloP or other measures of selection. When using this score, all significant association signals observed for the coding load score disappeared and no significant association for genome-wide and non-coding unweighted score was detected (S2 Fig). We further tested the associations with burden scores while restricting to only rare variants (DAF < 5%) or only common variants (5% < DAF < 70%, S2 Fig), however no significant association was observed. This is likely due to the domination of the mutation burden by alleles under effectively no purifying selection, highlighting the need for a weighting scheme to identify correlations to the relative per-individual fitness.

To assess whether the observed significant associations are sensitive to reference bias, we included as a covariate the number of non-reference sites per individual in our association testing for the top results in Table 2 (S6 Table). Association results were very consistent, suggesting that reference bias is not likely a confounder. Similarly, associations between the phenotypes and load score remain significant when restricted to variants at which reference alleles are the same as predicted ancestral alleles (S7 Table). We also re-computed load scores using phyloPNH scores, which are phyloP scores calculated without human reference genome [4], and obtained similar but slightly less significant results, with all the 27 phenotypes yielded p-value < 6.13×10^−4^ (S8 Table).

### Associations with coding load score are enriched for nominal associations with disease

Phenome wide association test results showed that no single disease is significantly associated with the load score (all P > 0.05 after accounting for multiple tests using Bonferroni correction). However, rather than the load score having a strong effect on a single disease, we hypothesized that the load score may have subtle effects on many diseases, leading to an excess of weak associations that do not individually reach statistical significance. To test this hypothesis, we compared the number of phenotypes nominally associated with the load score (p-value < 0.05 without multiple test correction) to a null distribution generated by random permutation of individual load score values (Methods). For this analysis, we restricted to associations with clinical phenotypes defined by phecodes. Out of 539 phecodes, 46, 24, and 27 phecodes (S9 Table) were found to be nominally associated with coding load score, non-coding load score, and genome-wide load score respectively. The number of nominally significant associations for coding load score is significantly larger than the expected number under the null model (P=0.005), supporting this hypothesis (Fig 3). However, this analysis was not statistically significant for the genome-wide load score and the non-coding load score (P>0.05) (S3 Fig), suggesting that diseases are largely correlated to the effect of variants in coding regions. We repeated the permutation analysis for the unweighted burden score as negative controls. As expected, enrichment of week association between burden scores and diseases are not statistically significant (S4 Fig).

**Figure 3.**
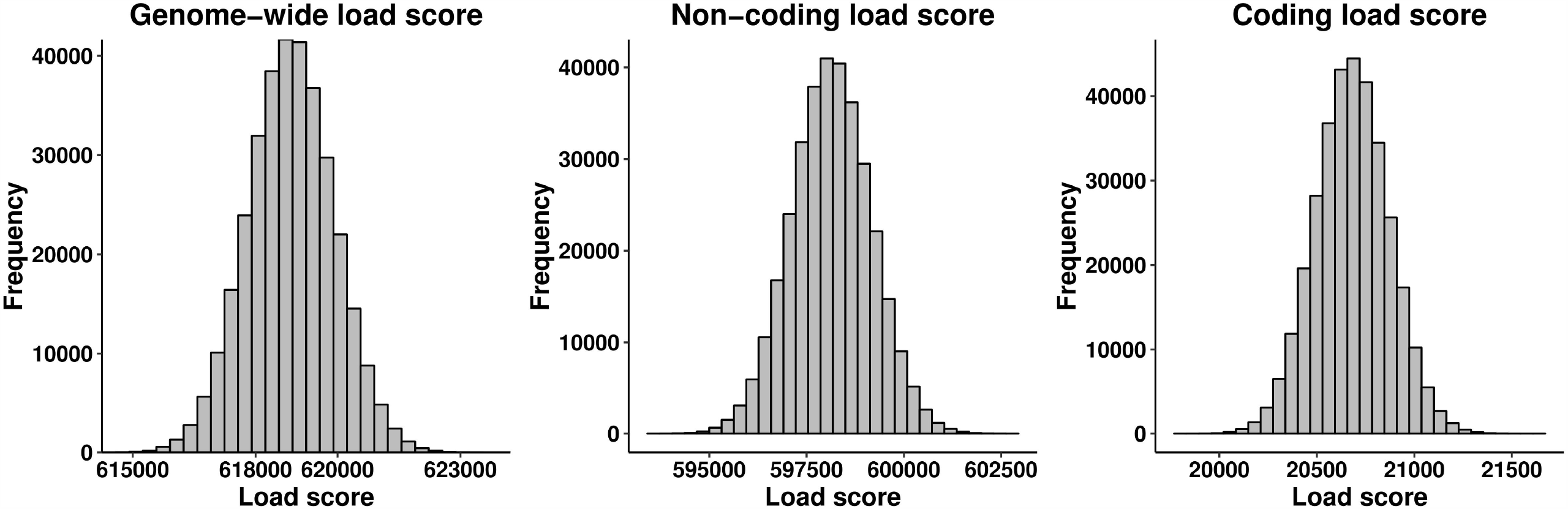
Enrichment of clinical phenotypes nominally associated with coding load score. Null distribution of the number of clinical phenotypes weakly associated with coding load score was obtained from 2,000 permutations in total. For each permutation, coding load score was shuffled randomly among 335,161 samples and the number of association was the count of phenotypes which yielded a p-value < 0.05 in the association tests between permuted load score and 539 phecodes. Red dashed line indicates the observed number of clinical phenotypes nominally associated with coding load score (n = 46).

## Discussion

In this study, we have described a polygenic load score that estimates the deleterious load carried by an individual, and applied this score to 335,161 white British individuals from the UK Biobank. Our analysis produced two major results: First, while we found no significant associations between individual medical phenotypes and the genome-wide load score, we found that more phenotypes are nominally associated with the coding load score than would be expected under a null model (Figure 3). This suggests that the deleterious load has a broad effect on the human phenome, rather than being specifically associated with a small number of phenotypes. This is consistent with Fisher’s Geometric Model of fitness, which proposes that the fitness of a population is determined by overall phenotypic distance from a theoretical optimal point in a phenotype space that potentially encompasses the organism’s entire phenome [29, 30]. Second, by restricting to protein coding variation, we found significant associations between the coding-only deleterious load score and a variety of adiposity phenotypes, along with other anthropometric phenotypes and phenotypes related to metabolic rate. This suggests that adiposity may be under polygenic selection driven by a large number of coding variants in humans. This is consistent with previous results obtained from the UK Biobank using an unrelated methodology [26]. We found no similar associations with the noncoding deleterious load score, which is in contrast to numerous studies finding significant genetic associations in noncoding regions, including associations with the same adiposity traits we found associated with our coding load score. Since our derived allele frequency spectrum analysis (Fig. 1) suggests that sites with higher phyloP scores are under purifying selection in noncoding regions as well as coding, the lack of significance in non-coding regions cannot be interpreted as a lack of purifying selection in these regions or poor sensitivity to selection in these regions. It may instead indicate that selection acts on phenotype associations in noncoding regions in a different way from how it acts in coding regions, possibly due to the small effect size of individual noncoding variants.

There are several limitations to our method. We computed the load score from imputed genotypes rather than sequenced whole genomes, which gives us little information about extremely rare variants in the population, masking potentially large contributions to the load from variants under the strongest selection. As a future topic of research, the same methodology can be applied to include rare variants, which would shed light on the relative contribution of common and rare variation to the phenotypic associations of load. Previous studies have shown that rare variation contributes substantially to differences in deleterious load between human populations, so we may expect it to have a significant impact on individual load in this context as well [1, 2].

Furthermore, the phyloP score used to estimate the deleteriousness of alleles measures only the likelihood that a site is evolving under constraint in vertebrates, and is not a direct estimate of the selective effect of a variant in humans. It is possible that the use of vertebrate-level conservation has reduced our ability to identify recent selection on human phenotypes, particularly those that are human specific. However, the fact that selection on adiposity traits was also detected by a method [26, 27] that does not rely on phyloP suggests that this result is not spurious. This feature of the phyloP score also makes it difficult to measure the effect of dominant or recessive selection, which may contain additional important insights. Finally, we did not incorporate any measure of positive selection in the computation of the load score. Scores similar to phyloP that could be used to detect positive selection do exist, but they rely on measures of nucleotide diversity and haplotype structure within larger regions of the genome, and are difficult to apply to single nucleotides as would be required to incorporate them into this analysis [31]. Methods to detect positive selection in the human lineage on finer scales are an area of ongoing research, and such methods could be incorporated into this approach in the future. All these methodological constraints limit the range of variation we identify as contributing to an individual’s burden score, and this limitation may be biased with respect to trait associations. In particular, we might expect rare variation or positive selection to reveal a different set of trait associations than the ones we find here by investigating common variation under purifying selection. This could potentially expand the scope of associated phenotypes beyond adiposity traits, a possibility that is also supported by the presence of nominal associations with many phenotypes unrelated to adiposity (S9 Table). Nevertheless, we do expect common variation under purifying selection to underlie a large fraction of common disease phenotypes, and therefore to provide valuable insights about the action of natural selection in humans.

One potentially exciting application for this approach is applying it to different populations to discover of population-specific insights into phenotypic associations with deleterious load. Since PhyloP scores can be calculated without any reference to specific human populations [4], there is no reason in principle that this method could not be applied to biobank data from other populations, given a sufficient number of samples. However, a few cautions are necessary. First, it is well known that comparisons of genetic associations and polygenic risk are unreliable across different ancestries [32], that signals of polygenic selection can easily be confounded by population structure or admixture [33, 34], and that mutation load specifically differs substantially between populations based on their demographic history [1, 2]. This makes it difficult to compare load scores directly between individuals of different ancestry, and also would likely make it difficult to apply this approach to admixed populations or populations with heterogeneous ancestry. Second, the approach of genotyping and imputation is entirely dependent on the availability of appropriate genotyping arrays and imputation panels, neither of which is necessarily available for all populations. It will be essential to use sequencing data for any population that is not well represented in these resources. Finally, many traits are strongly influenced by social, cultural, and environmental factors which may differ dramatically across populations, resulting in differences between populations that are not necessarily related to natural selection in a straightforward way. This is certainly true of the adiposity traits we identify in this study. Results of such studies should therefore be interpreted with caution.

The deleterious load score presented here provides a new approach to investigate the complex relationship between natural selection acting on individuals, individual medical phenotypes, and the human phenome at large. We expect that as the available biobank data continues to grow in size and scope, this method can be applied to larger and more diverse populations to gain additional insights into how load varies between different populations, possibly empowering population-specific medical discoveries with deleterious load.

## Material and Methods

### The dependence of derived allele frequency on deleteriousness score

We evaluated the dependence of derived allele frequency of single nucleotide polymorphisms (SNPs) discovered in the whole genome sequences of 7,509 non-Finnish European individuals in the GnomAD data set [23] on each of the four candidate annotations for the presence of purifying selection: GERP++ [15], phyloP [13], CADD [17], and fitcons [9]. 88,060,485 SNPs with less than one percent of missing data were considered. Functional effects and deleterious scores at each SNP were annotated using Whole Genome Sequence Annotator (WGSA) v0.7 [35]. We used functional effects annotated by Variant Effect Predictor (VEP) and determined derived and ancestral allele status based on the six-way EPO (Enredo, Pecan, Ortheus) multiple alignments of primate species.

For each deleteriousness score, we divided the SNPs into multiple groups with arbitrarily defined intervals based on the range of each score. The intervals used were: (0 ∼ 0.2, 0.2 ∼ 0.4, 0.4 ∼ 0.6, 0.6 ∼ 0.8) for fitcons, (−10 ∼ −7.5, −7.5 ∼ −5, −5 ∼ −2.5, −2.5 ∼ 0, 0 ∼ 2.5, 2.5 ∼ 5, 5 ∼ 7.5, 7.5 ∼ 10) for GERP, (0 ∼ 5, 5 ∼ 10, 10 ∼ 15, 15 ∼ 20, 20 ∼ 25, 25 ∼ 30, 30 ∼ 35) for CADD, and (−5 ∼ −2.5, −2.5 ∼ 0, 0 ∼ 2.5, 2.5 ∼ 5.0, 5.0 ∼ 7.5, 7.5∼ 10) for phyloP.

### Load score calculation

The load score of each individual was calculated by adding up the number (dosage in case of imputed SNPs) of derived alleles at each SNP, weighted by the phyloP score at that site, across the entire genome. Derived alleles were determined based on the six-way EPO alignment, as described above. Since we are focusing on the effect of purifying selection, only SNPs with positive phyloP score (positive scores denote uniform purifying selection, while negative scores denote clade-specific selection) were included. In this paper, we computed three load scores using three different SNP sets: the coding load score summed only over coding variants, the non-coding load score summed only over non-coding variants, and the genome-wide load score computed from both coding and non-coding variants. All load scores were computed using PRSice-2 software [36] under an additive model. Coding and noncoding variants were defined based on VEP annotation.

### Genotypic and phenotypic data

The UK Biobank consists of genotype, phenotype, and demographic data of more than 500,000 individuals recruited across the United Kingdom. Individual genotypes were generated from either the Affymetrix Axiom UK Biobank array (∼450,000 individuals) or the UK BiLEVE array (∼50,000 individuals), each contains ∼0.9 million markers. Additional variants were then imputed using the Haplotype Reference Consortium (HRC) combined with the UK10K haplotype resource, with a total of ∼96 million variants available in the latest released imputed data (version 3). To compute per-individual load scores, we restricted to variants with imputation quality INFO score >= 0.9. We excluded samples that were outliers in heterozygosity or missing rates, samples with putative sex chromosome aneuploidy, and samples with self-reported non-white British ancestry. We also excluded one individual from each pair of samples with relatedness up to the third degree. This produced a subsample of 335,161 individuals. All information used to exclude samples is included in the UK Biobank resource page.

UK Biobank provides a wide range of medical phenotypes from base line assessment, biochemical assays, dietary questionnaire, and health records. In the present study, we focused on 2,419 phenotypes which had been selected for heritability estimation by the Neale group (http://www.nealelab.is/blog/2017/9/15/heritability-of-2000-traits-and-disorders-in-the-uk-biobank). This subgroup covers phenotypes in most of the core categories, including early life and reproductive factors, family history, cognitive function, physical measures, lifestyle and health outcomes.

### Phenotype processing and association tests

Among the 2,419 phenotypes considered in our analysis, 619 phenotypes are international classification of disease (ICD-10) codes from electric health records. We converted ICD codes (including ICD-9 and ICD-10 codes) into phecodes using Phecode Maps 1.2 [37, 38]. This resulted in 1,677 unique phecodes in total. Of these, 539 phecodes with the number of cases greater than 500 were selected for phenome-wide association testing.

The remaining 1,800 phenotypes (2,419 – 619 ICD codes) were pre-processed using PHESANT [39], a package designed to process phenotypes and run phenome scans in UK Biobank. The PHESANT pipeline loads each input phenotype as continuous, integer, or categorical based on the information in the UK Biobank data dictionary; preprocesses and re-categorizes the phenotype data based on predefined rules; and assigns them into one of the four data types: continuous, ordered categorical, unordered categorical and binary. Of these 1,800 phenotypes, we only considered those with a minimum number of cases or controls equal to 500 and a minimum number of individuals equal to 5,000. This resulted in 841 phenotypes: 75 continuous, 104 ordered categorical, 36 unordered categorical, and 626 binary. In total, 1,380 phenotypes was included in our association analysis.

The association between load score and each phenotype was tested using a regression test in PHESANT: linear regression / lm R function for continuous, ordered logistic regression / polr R function for ordered categorical, multinomial logistic regression / multinom R function for unordered categorical, and binomial regression / glm R function with family = binomial for binary. Besides the commonly used covariates of age, sex, genotype chip, assessment center and 40 principal components, we added five variables as covariates in all association tests that might denote population structure: birth location, home area population density, Townsend deprivation index, and UK deprivation index.

### Association of 27 adiposity traits with coding load scores stratified by phyloP score and derived allele frequency

To explore which variants drive the association between the 27 adiposity traits and the coding load score, we stratified variants by derived allele frequency (rare variants, 0 to 0.05; intermediate frequency variants, 0.05 to 0.3; common variants, 0.3 to 0.7; and variants near fixation, 0.7 to 1) and phyloP score (0 to 2, 2 to 4, 4 to 6, 6 to 8, and 8 to 10). Simultaneous stratification was performed with four groups of SNPs: DAF<0.05 and phyloP≤2, DAF<0.05 and phyloP>2, DAF≥0.05 and phyloP≤2, DAF≥0.05 and phyloP>2.

### Permutation of phenome-wide association analysis and creation of null distribution

A null distribution of the number of clinical phenotypes weakly associated with load score was created by repeatedly running the association test between load scores and phenotypes after randomly shuffling the load scores of individuals within the tested sample. The phenotypes included in this permutation analysis were all 539 phecodes. The same set of covariates used in phenome-wide association study (PHEWAS) tests above was applied. For each permutation, the number of phenotypes nominally associated with the load score (p-value<0.05) was then computed. The permutation p-value was calculated as the fraction of permutations for which the number of nominally associated traits was at least as large as the observed number of nominally associated traits.

## Acknowledgments

This research has been conducted using the UK Biobank Resource under Application Number ‘16218’.

## Supporting information

**S1 Fig. Derived allele frequency spectra of coding and non-coding variants for different CADD and phyloP score categories**. Top row: Derived allele frequency spectrum of coding variants. Bottom row: Derived allele frequency spectrum of non-coding variants. Each solid line represents derived allele frequency spectrum of polymorphic sites belonging to one score category and three dashed lines represent derived allele frequency spectra of three control categories: synonymous (syn), missense (mis), and loss of function (LOF) variants.

**S2 Fig. Phenotypic association of load (phyloP-weighted) and burden (unweighted) scores**. Quantile-quantile plot of −log10 p-values for the phenotypic association of A) load scores (weighted by phyloP); B) burden scores (unweighted); C) burden scores restricted to rare variants (DAF<5%); and D) burden scores restricted to common variants (5%<=DAF<70%).

**S3 Fig: Enrichment of clinical phenotypes nominally associated with genome-wide load score and non-coding load score**. Null distribution of the number of clinical phenotypes weakly associated with genome-wide load score (left) and non-coding load score (right) was obtained from 2,000 permutations each. For each permutation, the load score was shuffled randomly among 335,161 samples and the number of associations on the x-axis was the count of phenotypes which yielded p-value < 0.05 in the association tests between the permuted load score and 539 phecodes. The red dashed lines indicates the observed number of clinical phenotypes nominally associated with genome-wide load score (n = 27, left) and non-coding load score (n = 24, right).

**S4 Fig: Enrichment of clinical phenotypes nominally associated with burden scores**. Null distributions of the number of clinical phenotypes weakly associated with burden scores were obtained using the same procedure to obtain the null distributions for load scores (Figure 3 and Figure S3). The red dashed lines indicates the observed number of clinical phenotypes nominally associated with genome-wide burden score (n = 22, left), coding burden score (n=20, middle), and non-coding load score (n = 20, right).

**S1 Table**. Linear regression between slopes and score categories.

**S2 Table**. Association between load score and the first 10 principal components.

**S3 Table**. Derived allele frequency stratification analysis.

**S4 Table**. phyloP score stratification analysis.

**S5 Table**. phyloP score and DAF stratification analysis.

**S6 Table**. Association between coding load score and the 27 phenotypes with number of non-reference variants included as a covariate.

**S7 Table**. Association between load scores restricted to sites where the human genome reference allele is the ancestral allele and 27 phenotypes.

**S8 Table**. Association between coding load scores computed from phyloPNH and 27 phenotypes.

**S9 Table**. Clinical phenotypes weekly associated with load scores.

