## Supplemental Table 1 for "Probing the aggregated effects of purifying selection per individual on 1,380 medical phenotypes in the UK biobank"

**S1 Table.** Linear regression between slopes and score categories.

| **Score** | **beta** | **SE** | **P-value** |
| --- | --- | --- | --- |
| fitCons | 0.061 | 0.012 | 0.037 |
| GERP | 0.094 | 0.021 | 0.0064 |
| CADD | 0.13 | 0.026 | 0.0046 |
| phyloP | 0.15 | 0.014 | 0.00039 |

Score categories from low to high are coded as an integer starting from 1. SE: standard error.
