## Supplementary figures and images for "Probing the aggregated effects of purifying selection per individual on 1,380 medical phenotypes in the UK biobank"

### Supplemental Figure 1

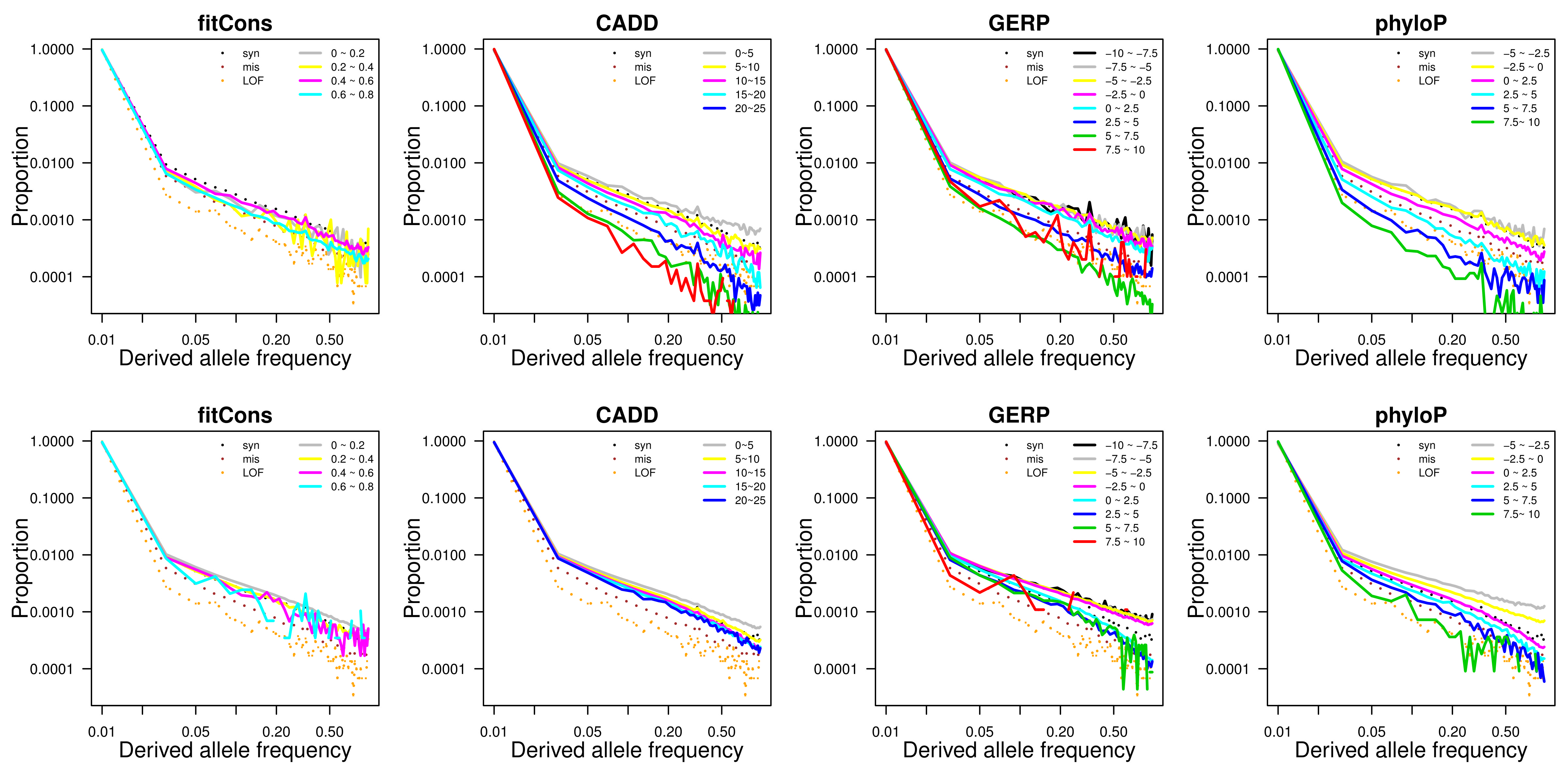

### Supplemental Figure 2

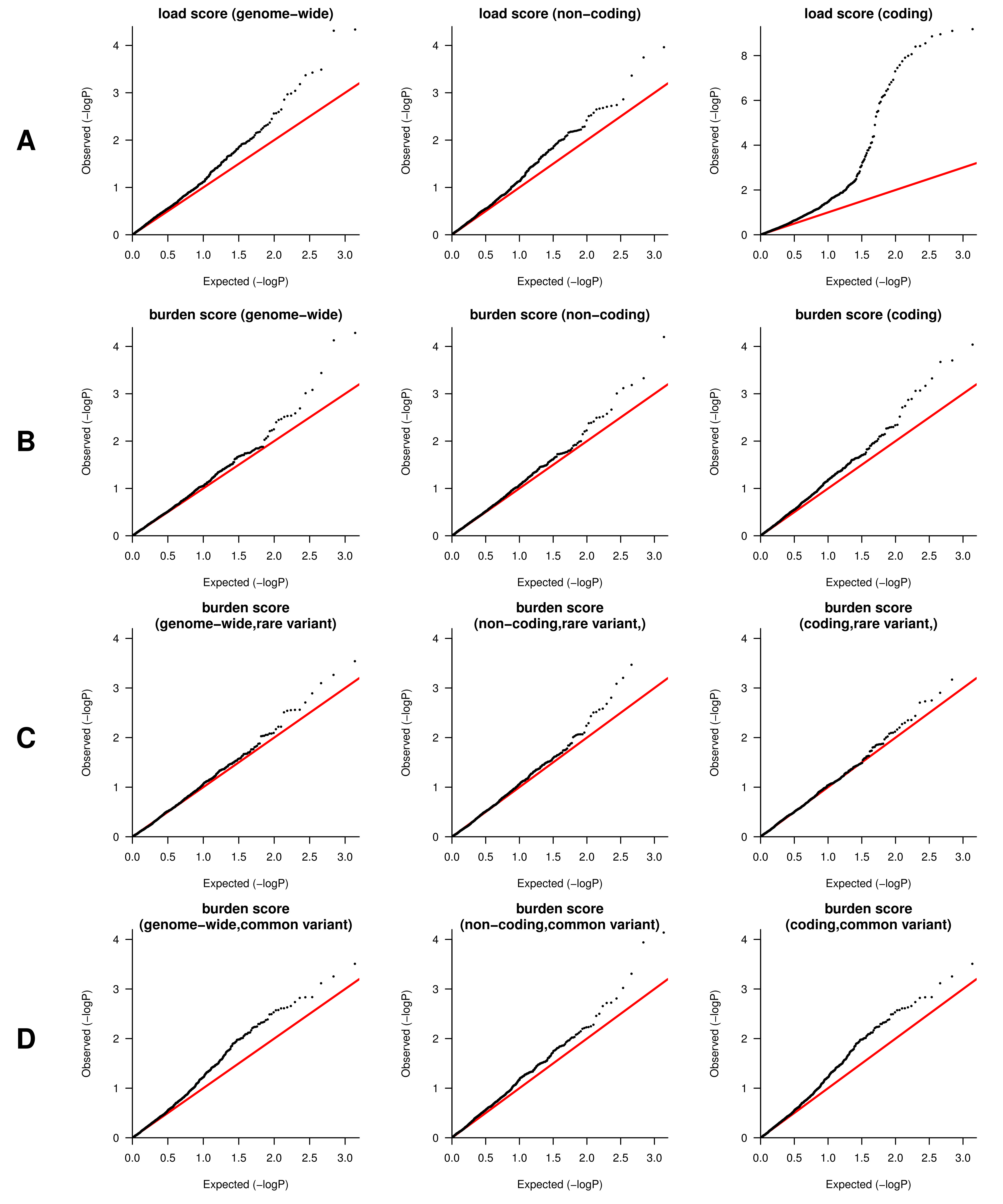

### Supplemental Figure 3

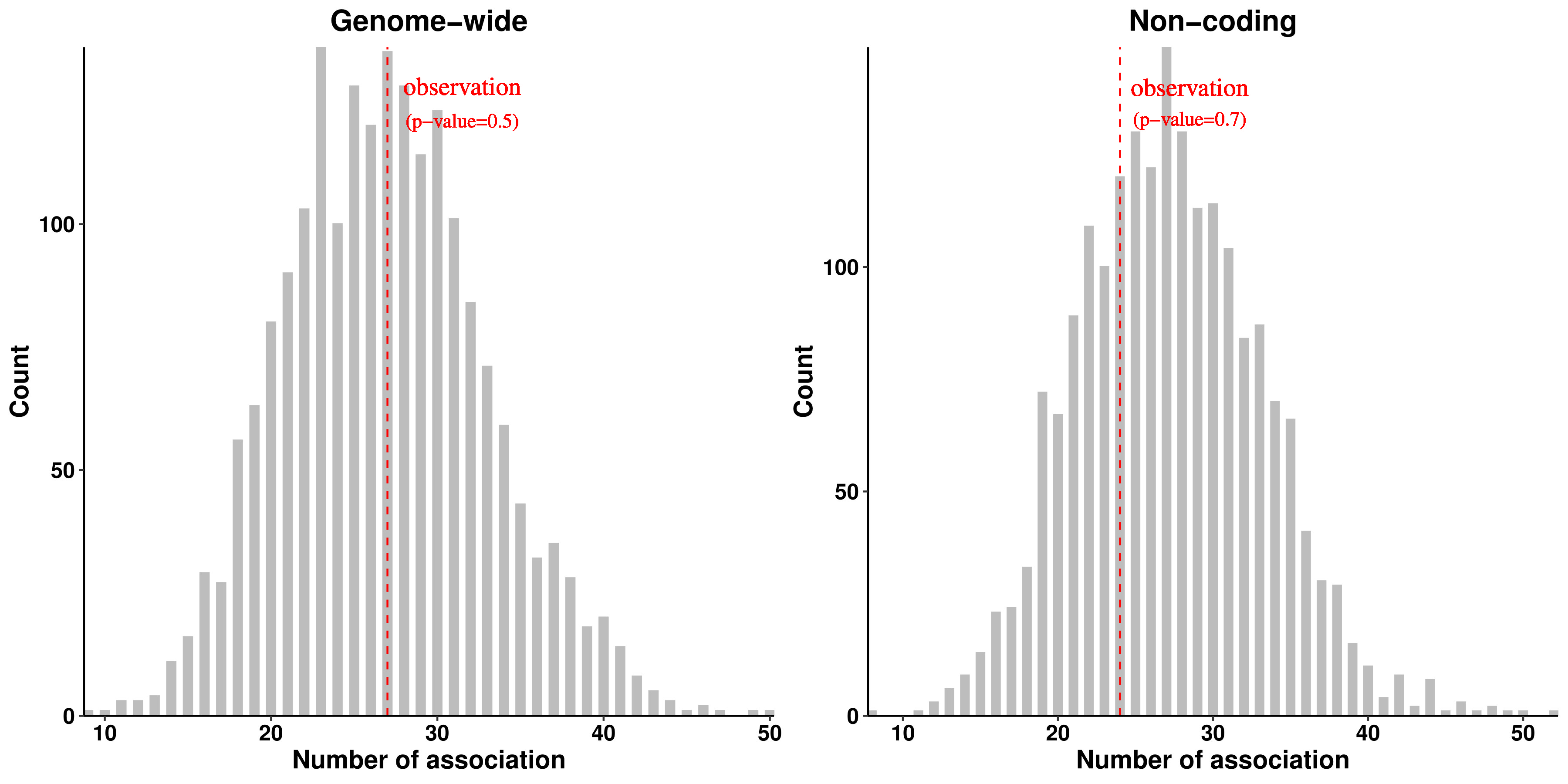

### Supplemental Figure 4

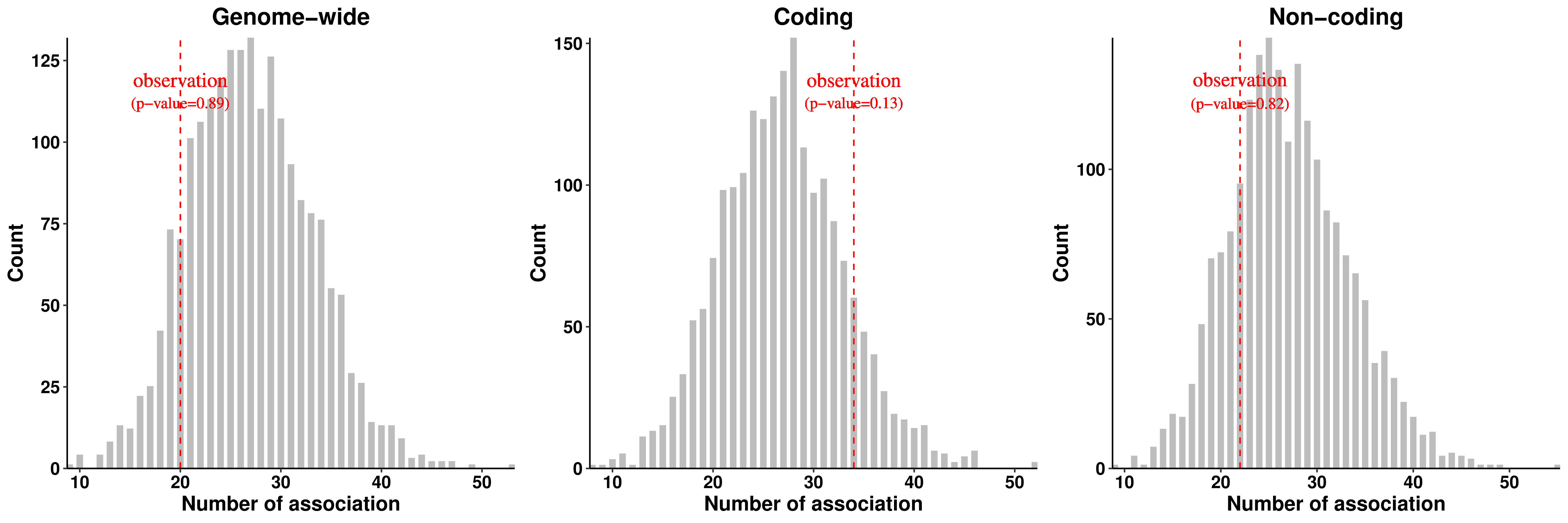
